# Variable effects on virulence of bacteriophage resistance mechanisms in extraintestinal pathogenic *Escherichia coli*

**DOI:** 10.1101/2022.09.01.506217

**Authors:** Baptiste Gaborieau, Raphaëlle Delattre, Sandrine Adiba, Olivier Clermont, Erick Denamur, Jean-Damien Ricard, Laurent Debarbieux

## Abstract

Bacteria exposed to killing agents such as antibiotics or viruses develop resistance. While phage therapy, the use of bacteriophages (phages) for treating bacterial infections, is proposed to answer the antibiotic resistance crisis, bacterial resistance to phages remains poorly characterized during phage treatment. We studied a large population of phage-resistant extra-intestinal pathogenic *Escherichia coli* 536 clones emerging from both *in vitro* (non-limited liquid medium) and *in vivo* (murine pneumonia) conditions. Genome sequencing revealed a mutational convergence of phage resistance mechanisms towards the modification of two cell-wall components, the K15 capsule and the LPS, whatever the condition, showing that their identification could be predicted from the *in vitro* conditions. The fitness cost of all phage resistant clones was broad in terms of growth rate and resistance to grazing by amoeba and could not discriminate K15 capsule to LPS mutants. By contrast, the virulence of the clones tested in mice showed that K15 capsule mutants were as virulent as the wildtype strain while LPS mutants were strongly attenuated. We also found that resistance to one phage led to the sensitization to other phages. In clinics, to control phage-resistant clones that remains virulent phage cocktail should include phages infecting both phage susceptible and future phage resistant clones.

**Importance:** *Escherichia coli* is a leading cause of life-threatening infections, including pneumonia acquired during ventilatory assistance for patients hospitalized in Intensive Care Unit, and a major multidrug resistant pathogen. A century-old concept, phage therapy (i.e. using specific anti-bacterial viruses), is being clinically re-evaluated supported with hundreds of successful compassionate phage treatments. However, along billions of years of coevolution bacteria have developed many ways to resist to phages. Phage resistance occurring during phage therapy remains often overlooked despite its critical role for a successful outcome. During this work we characterized phage resistant mutants in a virulent extra-intestinal pathogenic *E coli* strain and found that (1) phage resistance taking place during a phage treatment *in vivo* could be predicted from an *in vitro* assay; (2) phage resistance has, often but not always, a major fitness cost in terms of virulence; and (3) could be countered by appropriate cocktails of phages.

## Introduction

Antibiotic resistance constitutes a major threat for global public health, exposing patients to an increased risk of therapeutic impasse worldwide and increased mortality (1).

*Enterobacterales*, which include the genus *Escherichia*, are one of the most threatening multi drug resistant (MDR) bacterial families (1). In particular, *Escherichia coli* strains have become frequent etiological agents of ventilator-assisted pneumonia (VAP) (2, 3). Specific traits of extra-intestinal pathogenic strains of *E. coli* (ExPEC) involved in VAP include higher antimicrobial resistance and virulence factors, and belongs more often to ST127/B2 phylogroup (4, 5).

Within this context, phage therapy, the use of bacteriophages (phages), is increasingly considered for patients infected by MDR pathogens (6). Numerous experimental models and clinical compassionate-use case series support phage therapy efficacy (7–12). Remarkably, in experimental treatments a single dose of phages is often as efficient as multiple injections of antibiotics (13, 14). Nevertheless, these successes are currently offset by the lack of convincing clinical studies (15–17).

Like for antibiotics, bacteria have developed many ways to resist to phages (18). Intense research has revealed several novel phage-defense systems that target virtually all steps of the phage life cycle from adsorption to lysis (19, 20). Therefore, emergence of bacterial resistance to phages is possibly a critical factor, that could lead to treatment failure (21, 22). Interestingly, while phage resistance has been shown to develop rapidly *in vitro*, studies in humans have yielded discrepant results, reporting either the presence (23) or the absence (24) of phage resistance. This questions the importance of this phenomenon *in vivo*. Moreover, when phage resistance is studied *in vivo*, mechanisms involved are rarely investigated (25, 26).

A possible explanation for the lack of detection of phage resistance *in vivo* comes from the proposed synergistic action of phages with the immune system (27). Assaulted by two different antibacterial weapons, phages and immune cells, the density of bacteria rapidly decreases, which lowers the frequency of selection of phage-resistant clones. Alternatively, or complementarily, phage-resistant clones could display trade-off costs increasing their killing by immune cells. For instance, in several *in vivo* models of infection, resistance mechanisms resulted in altered virulence (14, 28, 29).

In order to gain mechanistic insights of phage resistance during phage therapy, we sequenced over 50 phage-resistant clones of the *E. coli* strain 536 exposed to the virulent phage 536_P1 collected from both *in vitro* (liquid medium) and *in vivo* (mice lungs) conditions. We previously reported that a single dose of this phage cures mice developing an otherwise fatal pneumonia by the *E. coli* strain 536 (30). Here, we identify a mutational convergence from the two conditions towards genes involved in either LPS or K15 capsule biosynthesis. Fitness studies of phage-resistant clones performed *in vitro* could not distinguish between the two classes of mutants. By contrast, a virulence assay in mice revealed that LPS-related mutants were almost avirulent while K15 capsule-related mutants were as virulent as the wildtype (WT) strain. Finally, we found that some of the phage-resistant clones could become susceptible to other phages that are unable to infect the WT strain.

## Results

### A single dose of phage 536_P1 selects phage-resistant clones more rapidly *in vitro* than *in vivo*

To assess the frequency of phage resistance emergence, as well as the molecular mechanisms involved, in two conditions, *in vitro* in non-limited liquid broth and *in vivo* during a phage therapy treatment of a murine pneumonia, we challenged the ExPEC strain 536 with a single dose of the virulent *Myoviridae* 536_P1 (30). We collected and re-isolated more than 90 clones per condition from multiple replicates (table 1). While less than 0.4% of the naïve population is resistant to phage 536_P1, the mean resistance rate is about 40% after 4 h of incubation in nutrient-rich liquid medium. In vivo, 13% of the clones recovered from mice lungs 10 h post-infection are resistant. This rate reached about 40% at 48 h (table 1). We observed that the bacterial resistance rate is heterogeneous among the replicates in all conditions suggesting that different resistance mechanisms could be involved (supplementary S1). To obtain a naïve population of bacteria exposed to the mice but not the phage, we also collected clones from the lungs of untreated mice 24 h post-infection and found that they all remained susceptible to phage 536_P1 (table 1).

### Genome sequencing reveals a mutational convergence of phage resistance mechanisms towards the modification of two cell-wall components

The genome of a subset of 57 phage-resistant clones from the above experiments was sequenced, revealing 65 mutations, including 53 different mutations, with at least one mutation per clone. By contrast, all phage-susceptible clones from control mice (*n* = 19) do not carry any mutation as compared to the original WT strain 536 (table 1; supplementary S2). Out of 53 unique mutations, 50 are expected to alter severely the function of the encoded proteins: 40 are truncating mutations and 10 are *in silico* predicted as deleterious mutations for protein function. None of them is specifically associated to the origin of samples (*in vitro* vs *in vivo*) (supplementary S2).

A large proportion (85%) of phage-resistant clones have at least one mutation in a gene involved in the LPS biosynthesis pathway, which overall include 77% of all mutations. The remaining mutations are located in the K15 capsule coding region (18%) and in the three following genes, all coding membrane proteins (5%): *mdtC* encoding an efflux system protein (31); *pqiA* involved in a membrane stability system (32); and *ECP_0298* predicted to be a homolog of a surface adhesin precursor (figure 1).

**Figure 1.**
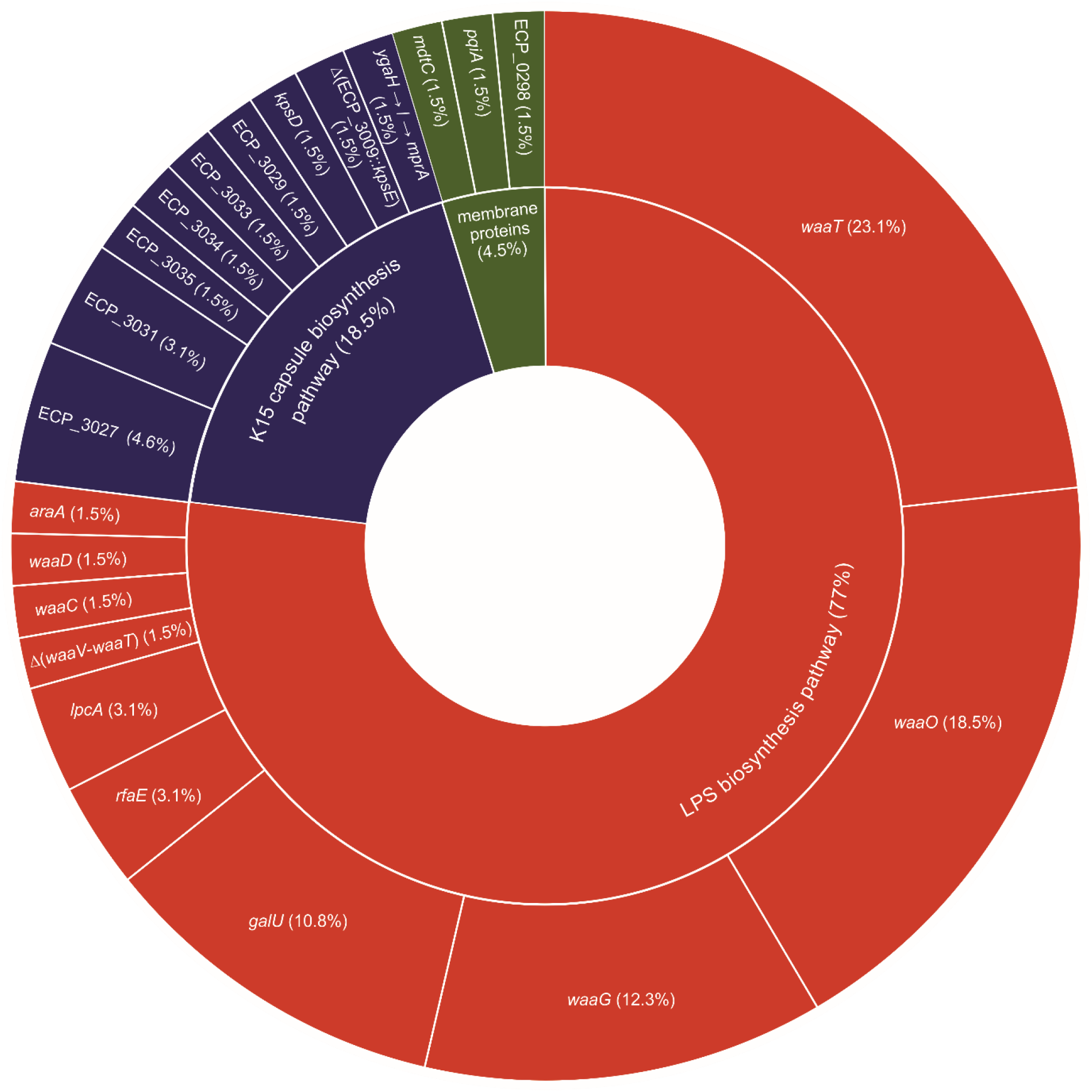
LPS and K15 capsule biosynthesis pathways are the two main targets of mutations identified from phage-resistant clones. Colours correspond to biosynthetic pathways. For each gene the percentage over the total number of mutations is indicated (*n*=57 clones).

By plotting the correspondence between mutated genes and the origin of samples (figure 2a) and by an MCA analysis (figure 2b), we found that none of the mutations is specifically associated with either *in vitro* or *in vivo* samples, showing that phage 536_P1 exerts a similar selection pressure on strain 536, independently of the environment where it infects it, liquid broth or mice lungs. The only obvious singularity are the three clones mutated in membrane proteins that originate from the lungs of infected and treated mice collected at 10 h post-treatment. However, these mutations are always associated with another mutation in a gene involved in LPS or K15 capsule biosynthesis. Nevertheless, while the same pathways (LPS and K15 capsule) are affected in both conditions, strictly identical mutations remain very low (three identical mutations among 65 different mutations were found in both conditions), showing a lack of hotspots (supplementary S2). Interestingly, the mutations located in LPS or K15 capsule related genes are mainly exclusive as only two out of 57 clones display mutations in both pathways (figure 2a).

**Figure 2.**
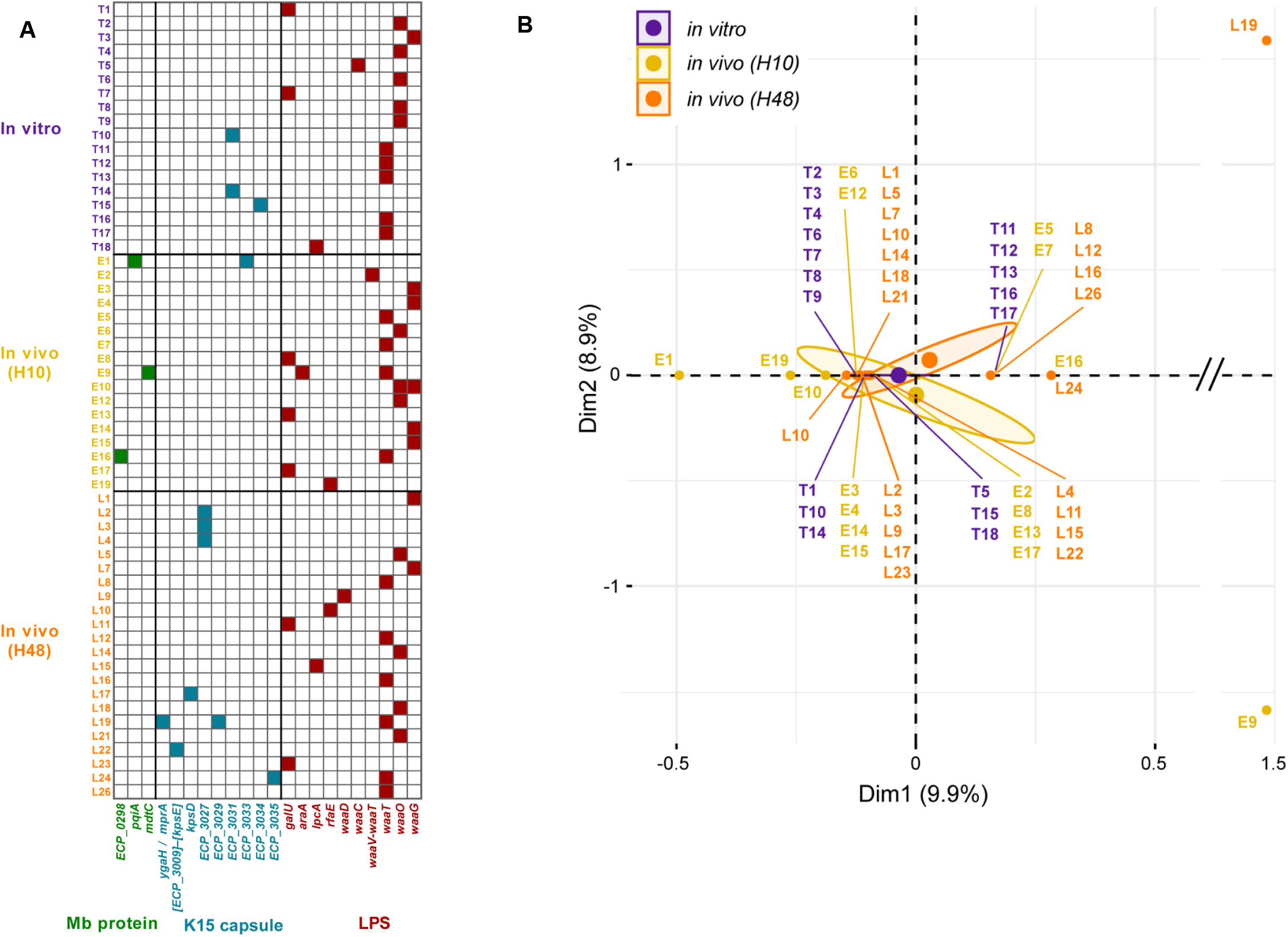
Genome sequencing of the 57 phage-resistant clones reveals a mutational convergence of phage resistance mechanisms at two levels (genes and metabolic pathways). **(A)** Mutated genes for each of the 57 clones are grouped per colour-coded conditions (*in vitro* and *in vivo* at 10 h and 48 h) and colour-coded metabolic pathways. **(B)** Multiple correspondence analysis computed according to mutated genes and metabolic pathways. *LPS*, lipopolysaccharide; ***Mb***, membrane.

These data argue for a mutational convergence whatever the condition, a hallmark of similar selection (33).

### The complementation of LPS but not K15 related mutants restore phage susceptibility

LPS is a surface exposed molecule well-known for its role in phage adsorption and many mutations in several genes involved in its biosynthetic pathways are associated to a loss of phage infection (34). The identified mutations would likely result in various degree of truncation of the LPS, as they occur either in the *waa* (or *galU*) cluster, which is involved in the assembly of the inner and outer cores, or in the *gmH* cluster (*lpcA, rfaE, waaD*), which is required for the synthesis of heptose (supplementary S3). To formally demonstrate that phage 536_P1 uses LPS as a receptor, we trans-complemented some LPS related mutants with a plasmid expressing the WT version of the mutated genes. Both phage 536_P1 susceptibility and adsorption were restored (supplementary S4a,b). By contrast, the same strategy applied to K15 capsule mutants failed to restore phage susceptibility (supplementary S4a), suggesting that instead of a loss of capsule synthesis these clones may produce a more abundant capsule to prevent phage infection. By using a K15 capsule specific serum we found that these mutants display a strong phenotype of agglutination by contrast to the WT strain 536 or a K15-negative control strain (supplementary S4c). Interestingly, none but one of the K15 capsule related mutations affects the capsule operon promoter (supplementary S2). We concluded that strain 536 deploys mainly two strategies to escape phage 536_P1 predation by modifying its receptor or making it inaccessible. As these two phage defence systems are nearly exclusive, we next evaluated the fitness cost of these mutations.

### The fitness cost of phage-resistant clones is dependant of the target of mutation and the environment

To determine the potential fitness cost of the various phage-resistant mutations, we studied their behaviour in three environments of increasing complexity.

First, the maximum growth rate (MGR) in non-limited nutrient medium of all clones with distinct mutations (*n*=47) was measured. Compared to WT strain 536 that has a MGR of 0.73 h^− 1^, clones mutated in LPS or in K15 capsule biosynthetic pathways have significantly lower MGR with a median of 0.53 h^− 1^ (range 0.33 to 0.68 h^− 1^; *p*=0.027) or 0.58 h^− 1^ (range 0.47 to 0.61 h^− 1^; *p*=0.042), respectively, while with a median MGR of 0.61 h^− 1^ (range 0.60 to 0.67 h^− 1^; *p*=0.349) the three clones mutated in membrane proteins remain similar to the WT strain 536 (figure 3a; supplementary S5).

**Figure 3.**
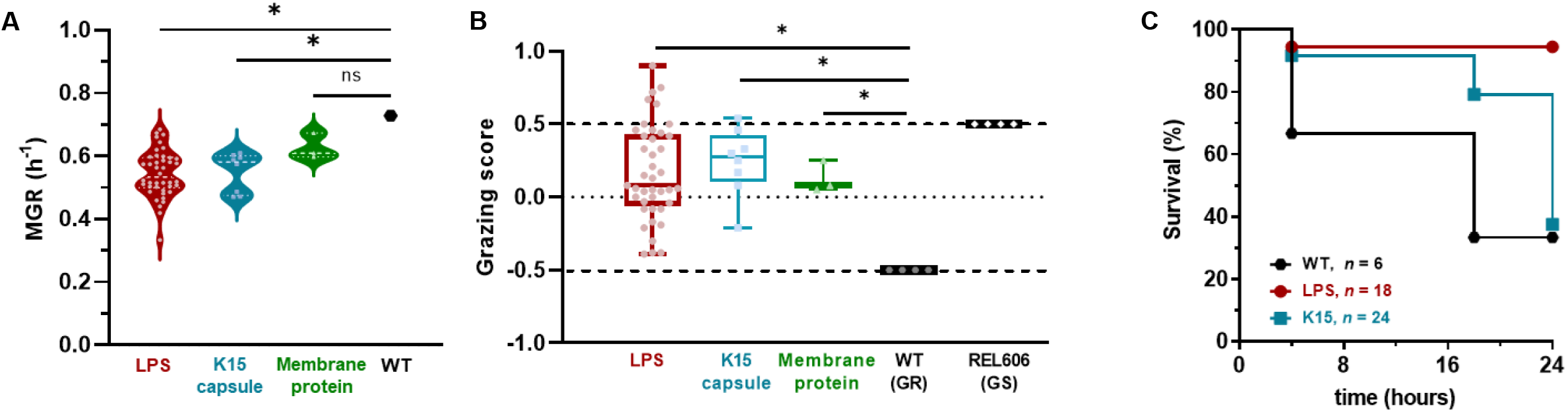
Phage resistant clones display a broad fitness cost *in vitro* and *in vivo*. **(A)** The growth rate of the 47 clones with distinct mutations in non-limited nutrient medium LB was calculated from kinetics recorded every 15 min in a microplate reader during 10 h (*n*=3 for each clone). **(B)** Protozoan predation by *D. discoideum* of the 47 clones with distinct mutations was evaluated by their grazing score (*n*=3 for each clone). **(C)** Virulence in acute pneumonia model was assessed towards a subset of clones carrying either single mutations in the LPS (*n*=3) or in K15 capsule locus (*n*=3). The survival of mice during 24 h post-infection (4 10^8^ CFU intra-nasal) by either the WT strain 536 (*n*=6) or clones (*n*=3) carrying single mutations in the LPS biosynthesis pathway (*n*=6 for each clone) or clones (*n*=3) carrying single mutations in the K15 capsule locus (*n*=8 per clone) is represented as Kaplan-Meier. LPS, lipopolysaccharide; MGR, maximum growth rate; GR, grazing resistance; GS, grazing sensitive

Second, we used *Dictyostelium discoideum*, a unicellular organism mimicking macrophage phagocytosis (35–37), to assess the grazing resistance (GR) phenotype of the above clones (*n*=47). The WT strain 536 is fully resistant to amoeba phagocytosis with no lysis plaque (GR phenotype) at densities of 10^2^, 10^3^, 10^4^ and 10^5^ amoebae per 10^8^ CFU of bacteria, whereas the control strain REL606 is grazing susceptible (GS phenotype) characterized by large lysis plaques and the formation of fruit bodies (supplementary S6). All the mutated clones showed an increased susceptibility to *D. discoideum*, compared to the WT strain 536, with some being even more susceptible than the GS control. However, no significant difference in the amplitude of the grazing phenotype was observed between LPS and K15 capsule mutants (*p*=0.80) (figure 3b).

Finally, we wondered if the virulence of phage-resistant clones towards the murine pneumonia model could be affected. We selected six mutated clones carrying single mutations either in the LPS (*galU*, clone T1; *waaG* clone E15; *waaT*, clone T11) or in the K15 capsule (*ECP_3033*, clone E1; *ECP_3027*, clone L3; Δ(*ECP_3009–kpsE*), clone L22). Kaplan-Meier survival curves of mice infected by these clones show a significant decrease of the virulence (*p* = 0.001) of the LPS mutated group of clones (*n*=18; 94% survival at 24 h post-infection), while K15 capsule mutated group of clones (*n*=24; 62.5% of survival at 24 h pi) remain as virulent (*p*=0.24), both compared to the WT strain 536 (*n*=6; 33% survival at 24 h pi) (figure 3c). When plotted for each mutated gene, results remain similar (supplementary S7b). As expected from the survival curves, the median level of CFUs in mice lungs at 24 h post-infection is 3-log lower for LPS mutants compared to both WT strain 536 and K15 capsule mutants (supplementary S7a). Therefore, while the above two *in vitro* assays do not discriminate any of the mutants, the *in vivo* virulence assay clearly segregates LPS to K15 capsule mutants.

Five phage resistant clones have two or three mutations in their genome. Four of them have a mutation in *waaT*, a gene involved in LPS outer core biosynthesis, associated with a gene coding either for a membrane protein or capsule K15. No difference in bacterial fitness was observed between single *waaT* mutants and double or triple mutants, (MGR, *p*=0.684; grazing score, *p*=0.301). Further investigations will be needed to establish the respective roles of these mutations.

### Resistance to one phage can lead to susceptibility to another one

The selection of K15 capsule phage-resistant clones that remain as virulent as the WT strain 536 could lead to phage treatment failure. We then asked whether other phages could readily infect these clones as a counter measure. We selected five virulent phages (536_P3, CLB_P2, LF73_P1, LF110_P3, DIJ07_P1) (supplementary S8) previously isolated and characterized in our laboratory for their ability to form plaques on a large collection of *E. coli* VAP strains (4). Amongst these five phages, two of them (LF110_P3 and DIJ07_P1) are unable to infect the original WT strain 536 (figure 4a). We tested these five phages for their ability to form plaques on 536_P1-resistant clones that have distinct mutations targeting capsule related genes (*n* = 7). We obtained a susceptibility coverage ranging from 0% (536_P3) to 100% (CLB_P2), showing that the overproduction of the K15 capsule does not provide a broad protection against many phages. Unexpectedly, phages LF110_P3 and DIJ07_P1 infect one and two, K15 mutants, respectively (figure 4b). Susceptibility to these two phages is also observed when testing LPS mutants, with a much larger extent for phage DIJ07_P1 that infects 39% of them (figure 4c). Therefore, when strain 536 exposed to phage 536_P1 responds by LPS or capsule modifications, these phage resistance mechanisms can be bypassed using additional phages, even if some of them are initially unable to infect the WT strain 536.

**Figure 4.**
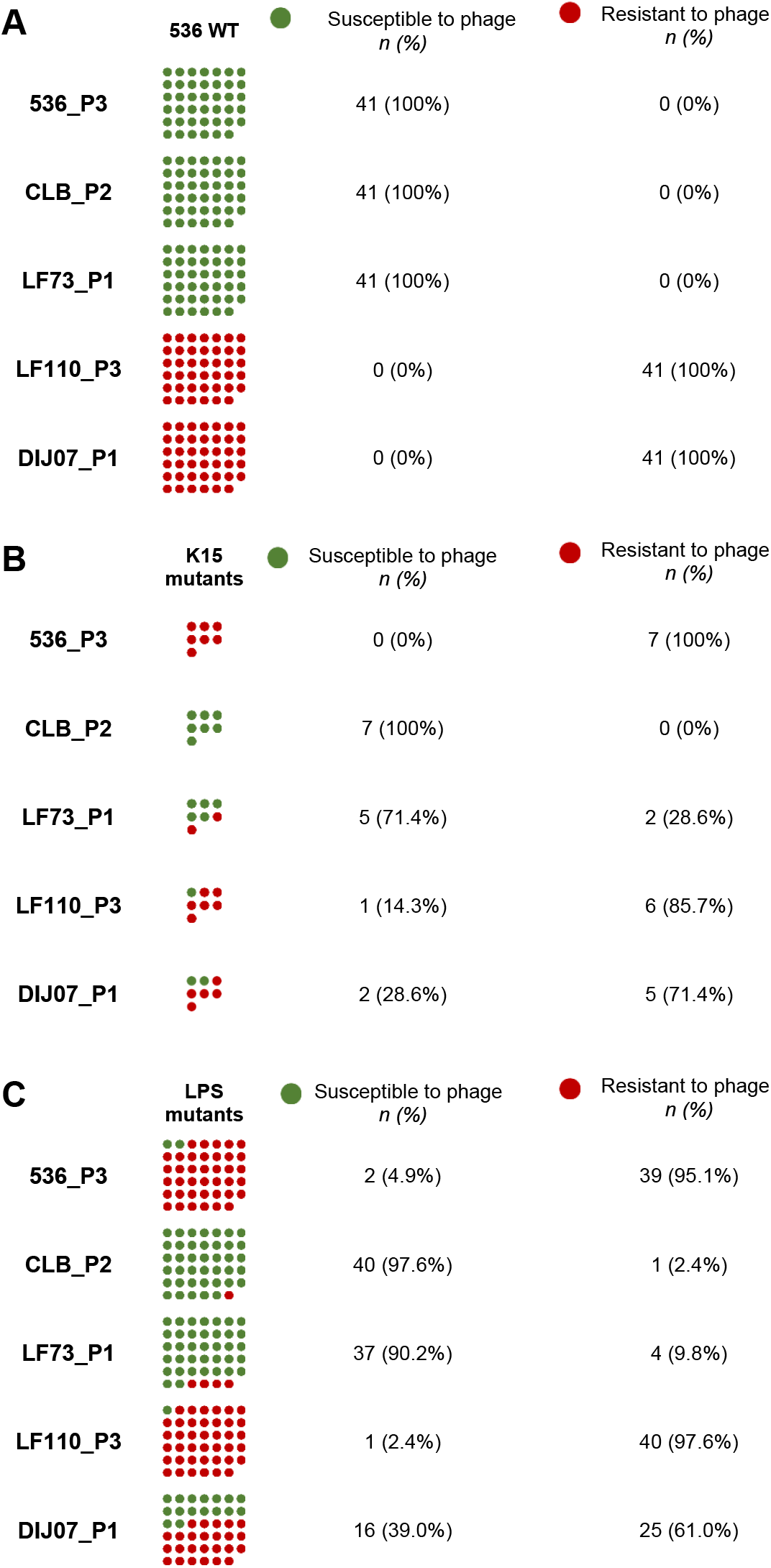
Phage 536_P1 resistant clones display uneven susceptibility to five other phages. Susceptibility tests of 41 clones of the WT strain 536 **(A)**, 7 phage 536_P1 resistant clones carrying a mutation in K15 capsule coding region **(B)** and 41 phage 536_P1 resistant clones carrying a mutation in a gene involved in LPS biosynthetic pathway **(C)** towards phages 536_P3, CLB_P2, LF73_P1, LF110_P3, DIJ07_P1.

## Discussion

Phage therapy has the potential to help patients infected by MDR bacteria (6). However, the current use of phages remains semi-empirical because of notably the lack of conclusive clinical trials. Moreover, the monitoring of phage resistance and the related mechanisms involved in patients undergoing treatment are often neglected although this resistance could be critical for the success of phage therapy (25, 26). Here, we studied a large population of phage-resistant clones from both *in vitro* and *in vivo* conditions and evaluated their mutational fitness costs. By contrast with several reports (14, 38, 39), phage resistance does not systematically drive the selection of less virulent clones but also selects for clones that remain as virulent as the WT strain, calling for a systematic monitoring of this phenomenon during treatment.

The sequence analysis of phage-resistant clones revealed a strong mutational convergence directed towards only two biosynthetic pathways, either the LPS or the K15 capsule. Nevertheless, the number of identical mutations remained rare, which argues for a lack of hotspots and support selection events taking place independently during each experiment. The two identified biosynthetic pathways have been previously reported as targets used by bacteria to defend themselves against phages *in vivo* (14, 28, 29, 40). Mechanisms involved in these two pathways of phage resistance are however different. LPS related mutations led the bacterium to interrupt phage receptor synthesis, while K15 capsule related mutants mask the receptor. With half of the bacterial genomes encoding at least one capsule biosynthetic cluster (41), including extra-intestinal virulent B2 phylogroup *E. coli* strains (42), the overproduction of capsule to provide phage resistance might be more frequent than anticipated. It is possible that the low number of phage resistant clones analyzed in former similar studies may have prevented the isolation of such mutants (26). We also cannot exclude that our observations may be restricted to this particular phage/bacterium couple. It is expected that phage resistance mechanisms affecting the bacterial envelope display a fitness cost compared to the WT phage-susceptible strain. This was indeed the case for all the mutants when measuring either their growth rates or their grazing resistance to amoeba. However, the lack of major fitness differences between LPS and K15 capsule related mutants towards amoeba remains surprising. Even if both LPS and capsule are envelope components involved in the pathogenicity of strain 536 (43–45), it was anticipated that LPS mutants would be more affected than capsule mutants. However, while serum resistance is directly linked to the presence of capsule in strain 536 (46) and contributes to grazing resistance (37), a previous study showed that mutants of strain 536 carrying an increasing number of deletions of pathogenicity islands did not become less resistant to grazing (47). The grazing test may therefore not be sensitive enough compared to the *in vivo* virulence assay, which revealed that capsule mutants were as virulent as the WT strain, while LPS mutant became avirulent. Therefore, the selection exerted by phage 536_P1 on strain 536 gives rise to two populations of resistant clones with opposite behaviors towards the animal host. However, since the frequency of capsule mutants is much lower than LPS mutants, the overall phage treatment remains successful (30). Nevertheless, we can reasonably expect that in other conditions, such as an immunodeficient context, the growth of these capsule mutants could lead to the failure of the treatment.

The time to reach 40% of mutants *in vivo* (48h) was much larger than *in vitro* (4h) most likely because of the time needed for the phage to reach and infect a large population of susceptible bacteria as well as the time needed for phage-resistant clones to grow. Nevertheless, in both conditions studied, the selection of phage-resistant clones led to mutations affecting the same biosynthetic pathways. This suggests that the animal host does not influence this selection process. However, the acute pneumonia model imposes limitations such as the use of a large bacterial population to initiate a fatal infection by overwhelming the immune system, as well as a large phage dose to provide a rapid and efficient treatment (48). It then remains possible that a putative impact of the animal host on phage resistance mechanisms could not been detected within this setting and may require extended time of observation. Nevertheless, we noticed that the frequency of K15 capsule mutants was higher at 48h compared to 10h post-infection, which is in agreement with their unaffected virulence. Then, the data presented here suggest that the characterization of phage resistant mutants selected *in vitro* could be sufficient to identify, at least, the main mechanisms by which bacteria will escape phage predation during the first phase of a treatment, in agreement with a recent compassionate treatment (49).

A counter measure to defeat the selection of phage-resistant mutants relies on the use of several phages, namely phage cocktails. However, the rules governing the choice of phages to be incorporated within a cocktail depends on the application. Currently, compassionate treatments include often several phages selected for their capacity to infect the strain of a patient, while commercial cocktails available in Georgia include phages infecting different bacterial species (50). Here, we found that some phage-resistant clones become susceptible to phages that cannot infect the WT strain 536. Similar phenomenon was recently reported (51). These observations suggest that phage cocktails may not necessarily need to include only phages infecting the patient’s targeted strain. Replacing phage therapy in a broader context of phage-bacteria coevolution could provide opportunities for each phage included in a cocktail to become at some point the most efficient phage (21).

In conclusion, we anticipate that the rapid analysis of phage-resistant mutants arising *in vitro* should be performed for each therapeutic phage in order to help building robust phage cocktails to be used in clinics.

## Materials and methods

### Bacterial strain, bacteriophage and culture conditions

*E. coli* strain 536 (4,938,920 bp; NC_008253.1) belongs to the B2 phylogenetic group, the O6:K15:H31 serotype and the sequence type 127 (52).

Phage 536_P1 (149.4 Kb) (30), was amplified on strain 536 and purified according to adapted molecular biology protocols (53) with an additional step of endotoxin removal using EndoTrap® (Lionex, Germany). The stock solution of 3.1 10^10^ PFU/mL (endotoxin concentration of 2.15 EU/mL) was diluted in phosphate buffered saline (PBS).

Unless otherwise specified, strains were cultivated at 37°C on LB medium (lysogeny broth) liquid or agar (Becton Dickinson, USA). Drigalski agar medium (Bio-Rad, France) was used to count bacteria from mice lungs. In amoeba experiments bacteria were cultivated in glucose-HL5 medium liquid or agar (Formedium, UK).

### Animals and Ethics statement

A total of 73 eight-week-old BALB/cJRj male mice (Janvier Labs, France) were housed in animal facilities in accordance with French and European regulations on the care and protection of laboratory animals. Food and drink were provided ad libitum. Protocols were approved by the veterinary staff of the Institut Pasteur (approval number 20.173) and the National Ethics Committee regulating animal experimentation (APAFIS#26874-2020081309052574 v1 and APAFIS#4947-2016020915474147 v5).

### Isolation of phage-resistant clones

#### From in vitro condition

A fresh exponential bacterial culture of a single *E. coli* strain 536 clone (*n*=7), from glycerol stock, was grown in LB medium under constant agitation at 37°C. Bacteriophage 536_P1 was added at MOI of 10^− 1^. The bacterial growth was monitored by measuring the turbidity using a spectrophotometer (Novaspec II, Pharmacia LKB, UK) at regular intervals. When the nadir was reached, remaining bacterial cells were centrifuged (4,000g, 15 min, 4°C) and the supernatant removed. The resuspended pellets were plated on LB agar plates and incubated overnight at 37°C.

#### From in vivo condition

We used a murine model of pulmonary phage therapy (30). Mice, anesthetized with a mixture of ketamine and xylazine by intra-peritoneal administration, were infected by intranasal inoculation with 1 × 10^8^ CFU of *E. coli* strain 536 (H0). Two hours later, mice, anesthetized by isoflurane inhalation (2%), were either treated by an intranasal administration of 20 μL of 536_P1 (i.e. 3 × 10^8^ PFU) (treated group, *n*=17), or either received an intranasal administration of PBS (control group, *n*=8). Mice were hydrated by subcutaneous route with 150 μL of 0.9% NaCl (Aguettant, France) and monitored twice a day to assess weight loss and behaviour. The treated group was split in an early follow-up group (H10, *n*=10) and a late follow-up group (H48, *n*=7). For the control group, mice were sacrificed at H24 by intraperitoneal injection of 150 μL of sodium pentobarbital (Ceva Santé Animale, France). Lungs were aseptically recovered, weighted and homogenized in PBS (FastPrep 25-5G, MP Biomedical, USA). Homogenates were centrifuged (4,000g, 15 min, 4°C), and the resuspended pellets in PBS were plated on Drigalski agar and incubated at 37°C.

### Determination of bacterial susceptibility to bacteriophage

From each animal and *in vitro* experiment, a maximum of 20 clones were randomly chosen and re-isolated three times on LB agar. The susceptibility to 536_P1 was assessed in liquid and solid media.

#### Growth kinetics in liquid medium

The bacterial growth was recorded by OD600nm readings every 15 minutes at 37°C under agitation for 10 hours using a microplate spectrophotometer (Tecan Infinite F200 pro, Switzerland). The OD600nm for each clone in two conditions (with and without phage) was measured in three different wells. The starting conditions were standardized: from a saturated culture, bacteria were refreshed to reach exponential growth from which the OD600nm was adjusted to 0.1, before to take 150 μL to fill a 96-well plate (i.e. 4.5 10^6^ CFU). Ten μL of bacteriophage (i.e. 1.5 10^6^ PFU) were added to experimental wells and 10 μL of PBS to control wells. A positive control (WT *E. coli* strain 536) was added in each microplate. The clones were classified into three categories according to their kinetic profiles: sensitive (equivalent to the WT *E. coli* strain 536 infected by 536_P1), resistant (equivalent to the WT *E. coli* strain 536 not infected), intermediate (any profile different from the two above).

#### Efficiency of plaquing (EOP) in solid medium

Bacteriophage 536_P1 was serially diluted (10 fold) in PBS and one drop of 4 μL of each dilution was spotted on LB agar plates previously inundated by a either the WT strain 536 or each mutant individually. The EOP was calculated as the ratio of the titre on a given mutant over the titre of the WT strain. A ratio of less than 0.5 indicates the presence of a resistant phenotype as previously defined (54).

### DNA extraction and Whole Genome Sequencing

Bacterial DNA extraction was carried out using the EZ1 DNA tissue kit (Qiagen, Germany) on EZ1 Advanced XL (Qiagen, Germany) according to the manufacturer’s recommendations. The libraries were prepared using the Nextera XT DNA Library Preparation Kit (Illumina, USA) according to the protocol previously described (55). A pair-end sequencing (300 bp) was performed on the MiSeq platform (Illumina, USA).

### Bioinformatics analysis of mutants

Reads quality was checked using FastQC (v0.11.8) (56). Reads were assembled with SPAdes (v3.11.1) (57). *In silico* typing of the clones was carried out using SRST2 (v0.2.0) allowing confirmation of both the sequence type (ST) according to the MultiLocus Sequence Typing (MLST) Pasteur (ST33) and Warwick (ST127), and the serotype (O6H31) (58). Phylogroup B2 of all phage-resistant clones was confirmed by the Clermont Typing software (59).

The WT strain 536 used for this study was sequenced together with phage-resistant clones. Compared to the referenced genome (NC_008253.1), some mutations were identified in the WT strain and integrated into the reference genome using gdtools of Breseq (v0.33). Mutations were identified by comparing the reads for each phage-resistant clone to the updated reference genome using Breseq Variant Report (V0.33) (60).

Each single nucleotide polymorphism detected was reviewed manually. For mutations located in genes of unknown function, protein-to-protein interactions search using the STRING database (61) was performed.

The functional consequence of mutations was predicted using SIFT (v1.3) (62), PolyPhen-2 (v2.2.2) (63) and PROVEAN (v1.1.3) (64). A mutation was considered potentially harmful if at least two of the predictions were concordant (EIPD score <0.05, a PolyPhen-2 result equal to “probably damaging” or “potentially damaging” and a PROVEAN score ≤ 2.5).

### Complementation of LPS and K15 capsule mutants

Phage-resistant clones carrying only one mutation in a gene involved in LPS biosynthetic pathway (*galU*, T7; *lpcA*, L15; *rfaE*, E19) were complemented by the corresponding ASKA plasmids (pCA24N backbone, chloramphenicol 25 μg/mL) (65) To complement K15 capsule-related phage-resistant clones (ECP_3022, L17; ECP_3031, T10) the corresponding ORFs were amplified from the WT strain 536 and cloned into the pUC18 plasmid (ampicillin 100 μg/mL) using *Bam*HI and *Sph*I enzymes (New England Biolabs, USA) (ECP_3022: F-GGATCCGAATGAGTTTGTGATGAAATTA, R-GCATGCTTACAAAGACAGAATCACTTTT; ECP_3031: F-GCATGCTTAAATTTCTGAGTACGGCAAA, R-GGATCCAATGGTGAAATATGAAAATCAA) Cloned ORFs were sequenced and plasmids transformed in the corresponding mutants.

### Phage adsorption

Adsorption of phage 536_P1 was assessed as described previously (66). Briefly, cells were infected at low MOI (0.1) and incubated 10 min at 37°C with constant agitation. 50 μL samples were collected every 30 sec until 7 min and kept on ice before titration on the WT strain 536 to determine the amount of free phages over time.

### K15 antiserum aggregation assay

Aggregation using a K:15 antiserum (Statens Serum Institut, Danmark) was performed as described (67). Briefly, 5 μL of two-fold serial dilutions of the K:15 antiserum (range dilution from 1:20 to 1:2560) were mixed with 10 μL (1.10^7^ CFU) of each bacterium tested and incubated at 37°C without agitation during 1 h and then at 4°C overnight.

### Fitness assays

#### Growth rate in non-limited liquid medium

For each strain, a single colony was picked from an agar plate to inoculate a single well of a 96-well plate filled with 150 μL of LB and incubated overnight under agitation at 37°C. 15 μL of each well was transferred into a new plate filled with 150 μL of LB and introduced into a microplate spectrophotometer to record OD600nm every 15 minutes at 37°C under regular agitation during 10 h. Three independent replicates were performed for each clone. The growth curves were analyzed with R software. For each curve, we normalized the initial time point as the first OD600nm measurement 0.1. We then smoothed the time series with the smooth.spline function and calculated the first (growth rate) and second derivatives, with respect to time, of the data expressed logarithmically. All time points at which the second derivatives changed sign (i.e., time points at which the growth rate was at a local maximum or minimum) were identified. We considered those to correspond to the maximum growth rate (MGR expressed in h^− 1^). Smoothing decreased the contribution of measurement noise to the maximum growth rate.

#### Dictyocellium discoideum grazing assay

The amoeba *D. discoideum* axenic strain AX3 and associated methods were described previously (37). Fresh bacterial exponential cultures, cultivated in 10 mL HL5 medium at 37°C under agitation, were washed in 10 mL of MCPB buffer, and adjusted to a bacterial concentration of 1.10^8^ CFU in 300 μL, which were plated on HL5 medium agar (55 mm diameter dishes). Fresh 24 h culture of *D. discoideum* cells grown in 10 mL of HL5 medium at 23°C with no agitation, were washed with 10 mL of MCPB buffer, and adjusted to five concentrations corresponding to 10, 10^2^, 10^3^, 10^4^, 10^5^ amoeba in 300 μL, which were overlaid on agar plates covered by bacterial lawns. Plates were incubated at 23°C and were examined at 3 and 6 days to record the appearance of lysis plaques, which translate the phagocytosis of the bacteria by the amoeba and defined the grazing sensitive (GS) phenotype, with strain *E. coli* REL606 used as a positive control. Lack of phagocytosis defines the grazing resistant (GR) phenotype knowing that the WT strain 536 was previously reported as GR. Results were expressed as a grazing score (number conditions with appearance of lysis plaques: 10^2^, 10^3^, 10^4^, 10^5^, and sporulation at D6). The scores were normalised for each independent experiment from −0.5 (GR of 536 WT) to 0.5 (GS of REL 606). Three independent replicates were performed by strain.

#### Virulence assay in mice

BALB/cJRj mice (*n*=48) were infected as described above and monitored to assess weight loss and behaviour three times within 24 h, at which point they were sacrificed to collect lungs that were processed as described above to count bacteria.

### Statistical analysis

Results are presented as the median and range, mean and standard deviation or the number and percentage of individuals.

Multiple Correspondence analysis (MCA) was performed with the mca function of the FactoMineR package (v 1.34) (68). MCA generated a simultaneous display of 57 observations (clones with at least one mutation) and 26 qualitative variables as condition with three modalities: *in vitro, in vivo* H10, *in vivo* H48; as LPS biosynthesis pathway or K15 capsule pathway or membrane protein coding gene, and as 22 different genes on a two-dimensional representation.

ANOVA was used to compare the MGRs in LB medium of the clones from three different conditions (*in vitro, in vivo* H10 and *in vivo* H48) and the WT strain 536, as well as grazing scores. All statistical analyses were carried out with GraphPad Prism software (v9.4.0) (GraphPad Software, USA) as well as Kaplan-Meier estimates of mouse survival. Survival differences were estimated by Log-Rank test. Significance is reached when the p value is <0.05.

## Acknowledgments

This work was partly funded by ANSM (Agence Nationale de Sécurité du Médicament et des Produits de Santé) HAP-2019-S003 to L.D., ANR (Agence Nationale de la Recherche) ANR-19-AMRB-0002 to JD.R. and L.D. and FRM (Fondation pour la Recherche Médicale) DEQ20161136698 to E.D. (équipe FRM 2016). B.G. received partial support from AIHP (Amicale des Anciens Internes des Hôpitaux de Paris). R.D. received support from Poste d’Accueil APHP/IP. We thank Hervé Le Nagard and Lionel de la Tribouille for the use of the CATIBioMed calculus facility and Thierry Pédron for his help with the agglutination assay.

