## Supplemental figures and tables for "Variable effects on virulence of bacteriophage resistance mechanisms in extraintestinal pathogenic *Escherichia coli*"

Supplemental information of the manuscript by Gaborieau et al. entitled:
Variable effects on virulence of bacteriophage resistance mechanisms in extraintestinal pathogenic *Escherichia coli*

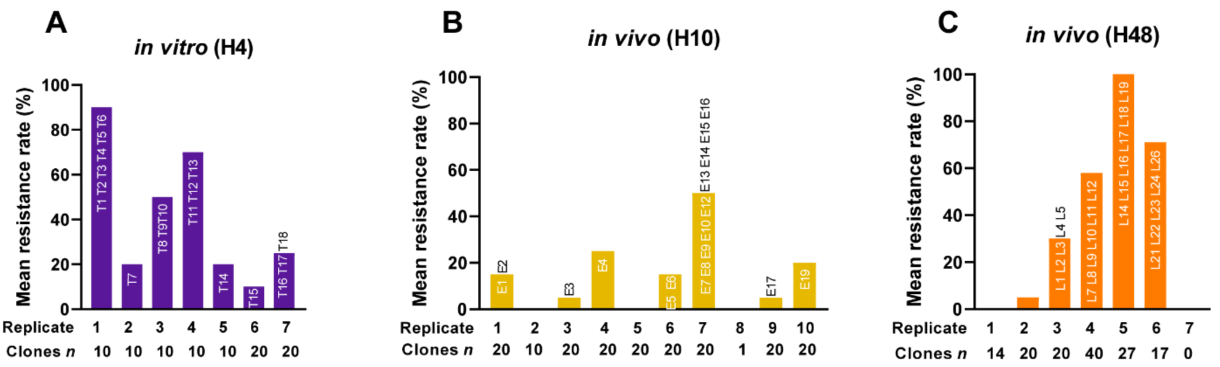


**Supplementary S1. Bacterial resistance rates to phage 536_P1 from the independent groups of both *in vitro* and *in vivo* conditions.**

**(A)** A total of seven liquid cultures of strain 536 were infected by phage 536_P1 at a phage:bacteria ratio of 1:10 during 4 h before sampling phage-resistant clones (*n*=10 to 20 per replicate).

**(B,C)** A total of 17 BALB/cJRj mice were infected by a single dose of 1 x $10^8$ CFU of strain 536 and treated 2 h later by a single administration of phage 536_P1 (3 x $10^8$ PFU). Mice lungs were collected 10 h (*n*=10) **(B)** and 48 h (*n*=7) **(C)** post-infection to sample phage-resistant clones (up to 20 clones per mouse). The identity of sequenced clones (*n*=57) reported in this study and listed in Supplementary S2 are indicated over the vertical bars.

| Clone[1] | Mutation type | Nucleotide change | Gene | Protein change | *In silico* prediction[2] SIFT | PROVEAN | POLYPHEN-2 | Function |
|---|---|---|---|---|---|---|---|---|
| **E16** | SNP | A→C | *ECP_0298* | Y160S | Tolerated (0.08) | Deleterious (-6.782) | Probably damaging (1.000) | Surface adhesin precursor |
| **E1** | SNP | T→C | *pqiA* | M314T | Affect protein function (0.04) | Deleterious (-4.524) | Probably damaging (0.998) | Membrane stability |
| **E9** | SNP | T→A | *mdtC* | L194Q | Affect protein function (0.00) | Deleterious (-5.815) | Probably damaging (1.000) | Efflux system |
| **L19** | SNP | A→G | *ygaH → / → mprA* | | | | | K15 capsule biosynthesis (transcriptionnal regulation) |
| **L22** | indel | del 8494bp | *[ECP_3009]–[kpsE]* | | | | | K15 capsule biosynthesis (export) |
| **L17** | SNP | G→T | *kpsD* | G32* | | | | K15 capsule biosynthesis (export) |
| **L2; L3; L4** | indel | del 1bp | *ECP_3027* | | | | | K15 capsule biosynthesis (export) |
| **L24** | indel | del 353bp | *ECP_3035* | | | | | K15 capsule biosynthesis (export) |
| **L19** | indel | del 1bp | *ECP_3029* | | | | | K15 capsule biosynthesis (synthesis) |
| **T14; T10** | indel | del 1bp | *ECP_3031* | | | | | K15 capsule biosynthesis (synthesis) |
| **E1** | indel | del 1bp | *ECP_3033* | | | | | K15 capsule biosynthesis (synthesis) |
| **T15** | SNP | A→T | *ECP_3034* | W281R | - | Deleterious (-7.875) | Probably damaging (1.000) | K15 capsule biosynthesis (synthesis) |
| **E9** | SNP | T→G | *araA* | Q287P | Affect protein function (0.01) | Deleterious (-5.997) | Probably damaging (0.998) | LPS biosynthesis |
| **T18; L15** | SNP | C→T | *lpcA* | P78L | Affect protein function (0.00) | Deleterious (-5.603) | Probably damaging (0.999) | LPS biosynthesis |
| **L23** | indel | del 1bp | *galU* | | | | | LPS biosynthesis |
| **E13** | indel | del 1bp | *galU* | | | | | LPS biosynthesis |
| **T1** | indel | del 1bp | *galU* | | | | | LPS biosynthesis |
| **E17** | indel | del 1bp | *galU* | | | | | LPS biosynthesis |
| **T7; L11** | SNP | C→T | *galU* | Q274* | | | | LPS biosynthesis |
| **L10** | indel | del 1bp | *rfaE* | | | | | LPS biosynthesis |
| **E19** | SNP | G→A | *rfaE* | T231I | Affect protein function (0.00) | Deleterious (-5.917) | Probably damaging (1.000) | LPS biosynthesis |
| **L9** | indel | del 1bp | *waaD* | | | | | LPS biosynthesis |
| **T5** | SNP | G→A | *waaC* | W47* | | | | LPS biosynthesis |
| **E2** | indel | del 2468 bp | *waaV–waaT* | | | | | LPS biosynthesis |
| **T17** | indel | del 1bp | *waaT* | | | | | LPS biosynthesis |
| **E7** | SNP | C→T | *waaT* | W290* | | | | LPS biosynthesis |
| **E5** | SNP | T→C | *waaT* | D212G | Affect protein function (0.00) | Deleterious (-6.863) | Probably damaging (1.000) | LPS biosynthesis |
| **L24** | indel | del 1bp | *waaT* | | | | | LPS biosynthesis |
| **L19** | indel | del 2bp | *waaT* | | | | | LPS biosynthesis |
| **E9** | indel | del 1bp | *waaT* | | | | | LPS biosynthesis |
| **T16; L12** | SNP | C→T | *waaT* | G177D | Affect protein function (0.00) | Deleterious (-6.916) | Probably damaging (1.000) | LPS biosynthesis |
| **T12** | SNP | C→A | *waaT* | E112* | | | | LPS biosynthesis |
| **L16** | SNP | G→T | *waaT* | Y105* | | | | LPS biosynthesis |
| **E16** | SNP | G→A | *waaT* | Q98* | | | | LPS biosynthesis |
| **T11; T13** | SNP | G→T | *waaT* | S80* | | | | LPS biosynthesis |
| **L26** | SNP | G→A | *waaT* | Q76* | | | | LPS biosynthesis |
| **L8** | indel | del 1bp | *waaT* | | | | | LPS biosynthesis |
| **T8; T9** | SNP | C→T | *waaO* | W294* | | | | LPS biosynthesis |
| **L21** | SNP | C→T | *waaO* | W276* | | | | LPS biosynthesis |
| **T2; T4; T6** | SNP | G→T | *waaO* | S252* | | | | LPS biosynthesis |
| **E6; E10; E12** | indel | del 1bp | *waaO* | | | | | LPS biosynthesis |
| **L5** | indel | del 1bp | *waaO* | | | | | LPS biosynthesis |
| **L14** | indel | del 1bp | *waaO* | | | | | LPS biosynthesis |
| **L18** | indel | +A | *waaO* | | | | | LPS biosynthesis |
| **L7** | indel | del 54bp | *waaG* | | | | | LPS biosynthesis |
| **E14; L1** | SNP | G→A | *waaG* | Q280* | | | | LPS biosynthesis |
| **L1** | SNP | G→A | *waaG* | R208C | Tolerated (0.06) | Deleterious (-4.221) | Probably damaging (1.000) | LPS biosynthesis |
| **E4** | indel | del 81 bp | *waaG* | | | | | LPS biosynthesis |
| **E15** | indel | del 18bp | *waaG* | | | | | LPS biosynthesis |
| **T3** | SNP | C→T | *waaG* | W71* | | | | LPS biosynthesis |
| **E3** | SNP | G→T | *waaG* | S58* | | | | LPS biosynthesis |
| **E8** | recombination | | *galU* | | | | | LPS biosynthesis |
| | | | *galU/hns* | | | | | LPS biosynthesis |
| **E10** | recombination | | *waaG* | | | | | LPS biosynthesis |
| | | | *waaG* | | | | | LPS biosynthesis |

**Supplementary S2. List of the 53 distinct mutations identified from the 57 phage-resistant clones sequenced.**

[1] T#, clones from *in vitro* condition; E#, clones from *in vivo* (10 h); L#, clones from *in vivo* (48 h).

[2] Predictions of the consequence of mutations on protein function.

| Phage-resistant clones | Mutated genes | Predicted LPS structure |
|---|---|---|
| 536 WT | none | 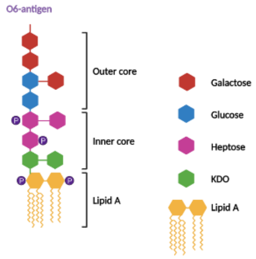 |
| T18. L15<br>E19. L10<br>L9 | *lpcA*<br>*rfaE*<br>*waaD* | 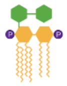 |
| T5 | *waaC* | 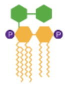 |
| T1. T7. E8. E13. E17. L11. L23<br><br>T3. E3. E4. E10. E14. E15. L1. L7 | *galU*<br><br>*waaG* | 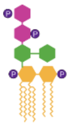 |
| T2. T4. T6. T8. T9. E6. E12. L5.<br>L14. L18. L21 | *waaO* | 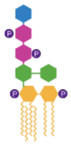 |
| T11. T12. T13. T16. E2. E5. E7.<br>E9. E16. L8. L12. L16. L19. L24.<br>L26 | *waaT* | 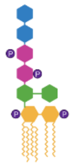 |

**Supplementary S3. Schematic representation of the LPS structures deduced from the mutated genes identified in phage-resistant clones.**

On the left column are indicated the list of the 47 clones carrying a point mutation in genes involved in the LPS biosynthesis pathway.

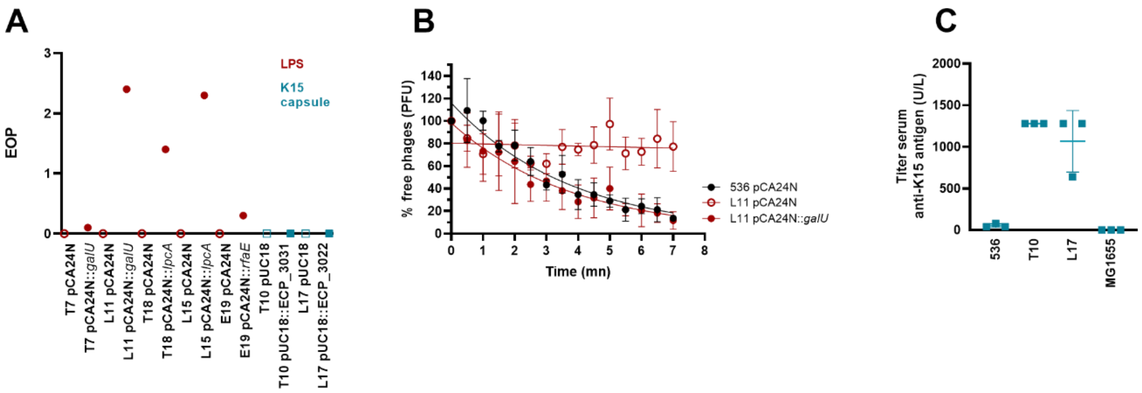


**Supplementary S4. Trans-complementation of LPS but not K15 capsule phage-resistant clones restore phage susceptibility.**

**(A)** Representative LPS (*galU*, clones T7 and L11; *lpcA,* clone T18 and L15; *rfaE,* clone E19) and K15 capsule (*ECP_3031*, clone T10; *ECP_3022*, clone L17) phage-resistant clones carrying a single mutation were trans-complemented by the corresponding WT gene cloned in a plasmid (pCA24N) and were assayed for their susceptibility to phage 536_P1. **(B)** Phage 536_P1 adsorption on the clone L11 (*galU*) is restored by trans-complementation. **(C)** Agglutination assays of two representative K15 capsule phage-resistant clones (L17 and T10) compared to strain 536 and the non-capsulated strain MG1655.

*EOP, efficiency of plaquing*

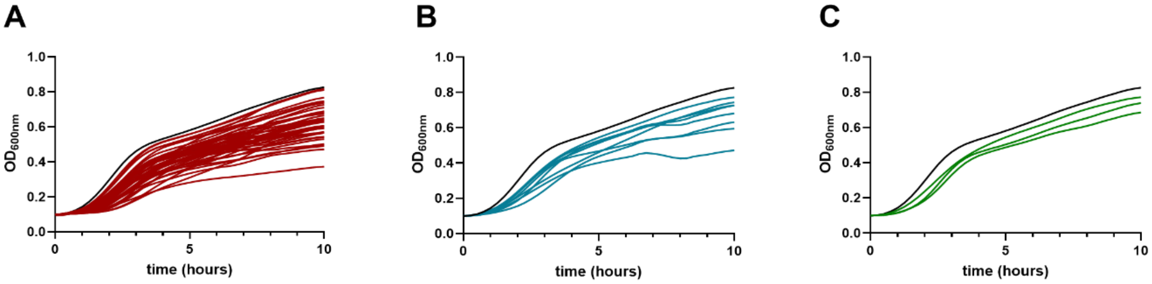


**Supplementary S5. The growth rate of phage-resistant mutants is lower than the wild-type strain 536.**

The growth of all sequenced phage-resistant clones with no identical mutation (n=47) in a non-limited nutrient medium (LB medium) is represented as the mean and smoothed growth curve of three independent replicates compared to the WT strain 536 (black) and displayed by mutated functions in **(A)** clones involved in LPS biosynthetic pathway (red), **(B)** mutated clones involved in K15 capsule genes (blue) and **(C)** mutated clones in membrane proteins (green).

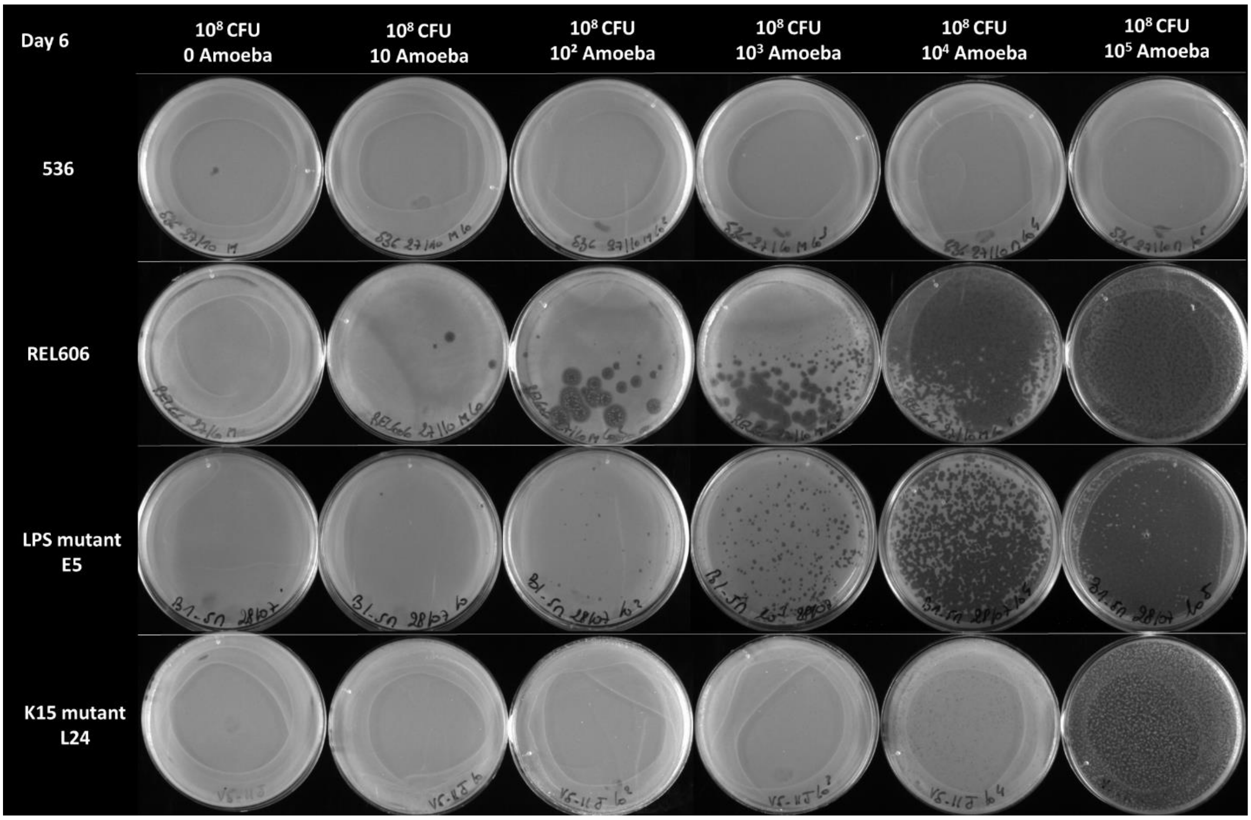


**Supplementary S6. Phage resistant clones become susceptible to the protozoan predation by *Dictyostelium discoideum*.**

Representative images of grazing resistant (strain 536) and grazing susceptible (strain REL606) phenotypes estimated by the serial dilution of amoeba cells from 0 (left) to $10^5$ (right) on bacteria lawns ($10^8$ CFU) as well as two phage-resistant clones (E5 and L24).

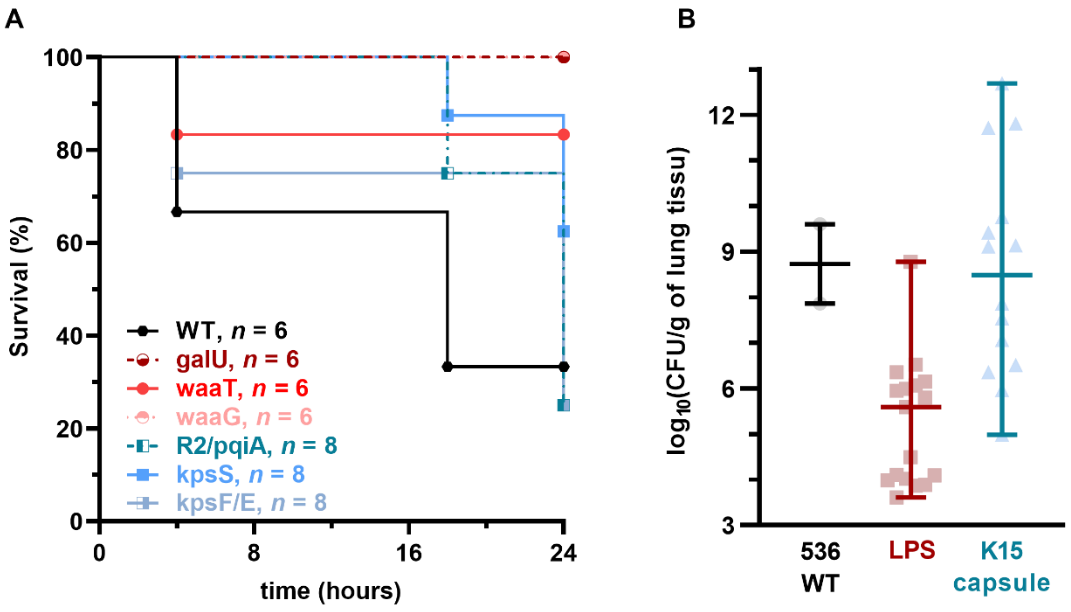


**Supplementary S7. The virulence of LPS but not K15 capsule phage-resistant mutants is strongly attenuated.** **(A)** Three representative of LPS mutants (*galU*, clone T1; *waaT*, clone T11; *waaG*, clone E15) and three K15 capsule mutants (*ECP_3033*, clone E1; *ECP_3027*, clone L3; Δ*(ECP_3009–kpsE)*, clone L22) were used to infect BALB/cJRj mice (a single dose of 1 x $10^8$ CFU; n=6 to 8 per group), whose survival rates over 24 h were monitored. **(B)** Lungs from infected mice described in **(A)** were collected 24 h post-infection to count bacteria.

**Supplementary S8.** List of the six phages used in this study.

| Bacteriophage | Morphology | Genus | Genome size (kb) | Infects WT strain 536 |
|---|---|---|---|---|
| 536_P1 | *Myoviridae* | *Phapecoctavirus* | 149.4 | Yes |
| 536_P3 | *Podoviridae* | *Phapecoctavirus* | 39.7 | Yes |
| CLB_P2 | *Myoviridae* | *Dhakavirus* | 171.8 | Yes |
| LF73_P1 | *Myoviridae* | *Tequatrovirus* | 168.9 | Yes |
| LF110_P3 | *Myoviridae* | *Felixounavirus* | 88.2 | No |
| DIJ07_P1 | *Myoviridae* | *Phapecoctavirus* | 142.8 | No |
